# Identifying gene targets for brain-related traits using transcriptomic and methylomic data from blood

**DOI:** 10.1101/274472

**Authors:** Ting Qi, Yang Wu, Jian Zeng, Futao Zhang, Angli Xue, Longda Jiang, Zhihong Zhu, Kathryn Kemper, Loic Yengo, Zhili Zheng, eQTLGen Consortium, Riccardo E. Marioni, Grant W. Montgomery, Ian J. Deary, Naomi R. Wray, Peter M. Visscher, Allan F. McRae, Jian Yang

**Author notes:** Correspondence: Jian Yang.

## Abstract

Understanding the difference in genetic regulation of gene expression between brain and blood is important for discovering genes associated with brain-related traits and disorders. Here, we estimate the correlation of genetic effects at the top associated *cis*-expression (cis-eQTLs or cis-mQTLs) between brain and blood for genes expressed (or CpG sites methylated) in both tissues, while accounting for errors in their estimated effects (*r*_*b*_). Using publicly available data (*n* = 72 to 1,366), we find that the genetic effects of cis-eQTLs (*P*_eQTL_ < 5×10^−8^) or mQTLs (*P*_mQTL_ < 1×10^−10^) are highly correlated between independent brain and blood samples (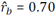 with SE = 0.015 for cis-eQTL and 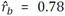 with SE = 0.006 for cis-mQTLs). Using meta-analyzed brain eQTL/mQTL data (*n* = 526 to 1,194), we identify 61 genes and 167 DNA methylation (DNAm) sites associated with 4 brain-related traits and disorders. Most of these associations are a subset of the discoveries (97 genes and 295 DNAm sites) using data from blood with larger sample sizes (*n* = 1,980 to 14,115). We further find that cis-eQTLs with tissue-specific effects are approximately uniformly distributed across all the functional annotation categories, and that mean difference in gene expression level between brain and blood is almost independent of the difference in the corresponding cis-eQTL effect. Our results demonstrate the gain of power in gene discovery for brain-related phenotypes using blood cis-eQTL or cis-mQTL data with large sample sizes.

## INTRODUCTION

Genome-wide association studies (GWAS) have discovered thousands of genetic variants associated with complex traits and diseases^1-3^. Most trait-associated variants reside in non-coding regions of the genome^4,5^, suggesting that genetic variants may affect the trait through regulation of gene expression^6,7^. With the advances in microarray and sequencing technologies, genome-wide genotype and gene expression data available from relatively large samples have been generated to identify genetic variants affecting transcription abundance^8-10^, i.e. expression Quantitative Trait Loci (eQTLs). Current eQTL studies are biased toward the most accessible tissues (e.g. blood), which are often not the most relevant tissues to the traits and diseases of interest. The Genotype-Tissue Expression (GTEx) project^11-13^ provides a comprehensive resource of data to investigate the genetic variation of gene expression across a broad range of tissues and cell types. Recent studies have utilized the GTEx data to demonstrate that the genetic correlation of gene expression between tissues in local regions (i.e. ±1Mb of the transcription start site) is much higher than that in distal regions^14^, consistent with the conclusions from the latest GTEx release^13^, and that there is no evidence for the tissue-relevant eQTLs being enriched for associations with complex traits^15^.

For studies that integrate GWAS results with eQTL or methylation QTL (mQTL) data to identify putative functional genes and regulatory elements for brain-related phenotypes and diseases^16,17^, the statistical power is limited by the small sample sizes of the brain eQTL or mQTL data (often in the order of 100s). On the other hand, there are blood eQTL and mQTL data available from thousands of individuals^8,9^ and the sample sizes of some of the ongoing projects have reached 10,000s (e.g. the GoDMC and eQTLGen consortia). The questions are to what extent the cis-genetic effects on gene expression and DNA methylation (DNAm) in blood differ from those in brain and whether we can gain power for detecting associations of genes (or DNAm sites) with brain-related traits by using the cis-eQTL (or cis-mQTL) effects estimated from a large blood sample as proxies for those in brain. In this study, we use a summary-data-based method to estimate the correlation of effect sizes of cis-eQTLs (or cis-mQTLs) between blood and brain for genes expressed (or CpG sites methylated) in both tissues, accounting for errors in their estimated effects. We then test whether there is an enrichment of the cis-eQTLs or cis-mQTLs with tissue-specific effects between blood and brain in the epigenomic states annotated by the ENCODE project^18^ and the Roadmap Epigenomics Mapping Consortium (REMC)^19^. We further implement a method (<u>m</u>eta-analysis of <u>e</u>QTL data from <u>c</u>orrelated <u>s</u>amples, MeCS) to meta-analyze cis-eQTL summary data from all the GTEx brain regions to maximize the power of detecting eQTLs in brain. We demonstrate by simulation and analysis of real data the gain of power by using cis-eQTL or cis-mQTL effects estimated in blood as proxies of those in brain to identify putative functional genes for brain-related complex traits and diseases. Almost all the analyses were performed based on summary-level data from previous studies.

## RESULTS

### Estimating the correlation of cis-eQTL effects between brain and blood

To quantify the similarity of genetic effects at the top associated cis-eQTLs (or cis-mQTLs) between two tissues, we used a summary-data-based approach to estimate the correlation of cis-effects between two tissues (*r*_*b*_) correcting for errors in the estimated cis-eQTL (or cis-mQTL) effects and sample overlap (**Supplementary Fig. 1 and Methods**). We show by simulation (**Supplementary Note**) that *r*_*b*_ is a good estimator of the correlation of the true values of cis-genetic effects (**Supplementary Fig. 2**). Note that the *r*_*b*_ method is distinct from the Spearman or Pearson correlation approach^13^ because the latter does not account for errors in the estimated eQTL effects and thereby leads to an underestimation of the correlation of true eQTL effects. We applied our method to estimate 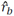 at the top cis-eQTLs between different brain regions and between brain and blood in one data set, and between brain and blood in two data sets using summary-level data from GTEx v6 (whole blood and 10 brain regions)^11^, the CommonMind Consortium (CMC; dorsolateral prefrontal cortex)^20^, the Religious Orders Study and Memory and Aging Project (ROSMAP)^21^, and the Brain eQTL Almanac project (Braineac; 10 brain regions)^22^ (**Methods and Supplementary Table 1**). All eQTL effects were in standard deviation (SD) units. For the GTEx, CMC and ROSMAP data, which are based on RNA sequencing (RNA-Seq), we matched the data sets by Ensembl Gene IDs. For the Braineac data that are based on gene expression microarray, we matched the data sets by gene symbols and removed genes tagged by multiple gene expression probes to ensure a one-to-one match for genes between data sets. The main aim of our study is to quantify the extent to which cis-eQTL data in blood can be used for the identification of genes associated with brain-related phenotypes and disorders. However, if we had selected eQTLs as the most associated SNPs in a linkage disequilibrium (LD) region for one tissue (say blood) and compared their effects with those in the other tissues (say brain), we would likely suffer a form of winner’s curse. To avoid potential ascertainment bias, we selected the top cis-eQTLs in a reference tissue, i.e. GTEx-muscle (*n* = 361) or CMC (*n* = 467; independent of GTEx), and estimated *r*_b_ between brain and blood. We used a stringent p-value threshold that is required for the SMR analysis^23^ (see below) to select cis-eQTLs. Then, we estimated the *r*_b_ between two tissues using these SNPs (**Supplementary Fig. 3**). Although this strategy uses only a quarter of all genes, the estimates of *r*_*b*_ should be valid (see below).

**Figure 1.**
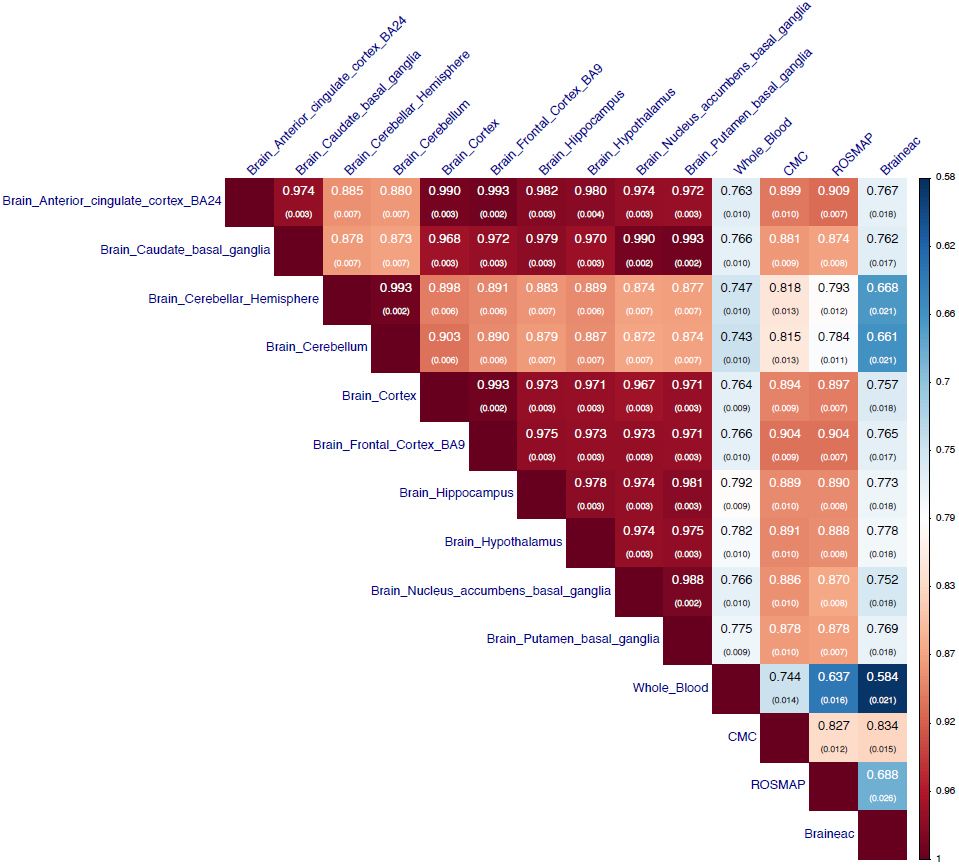
Estimated correlation of genetic effects of cis-eQTLs between brain regions, between brain and blood tissues, and between data sets. The top associated cis-eQTLs (one for each gene) were selected from GTEx-muscle at *P*_eQTL_ < 5×10^−8^. Shown in each cell is the estimate of *r*_b_ with its standard error given in the parentheses (**Methods**). In the Braineac data, the eQTLs effect sizes were estimated from gene expression levels averaged across 10 brain regions.

**Figure 2.**
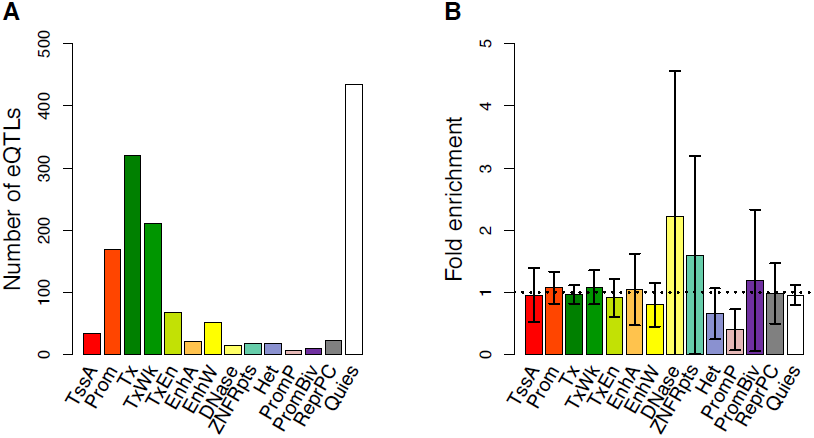
Enrichment of cis-eQTLs with tissue-specific effects in functional annotations. A) The distribution of cis-eQTLs across 14 functional categories derived from RMEC (**Methods**). B) Estimated enrichment of *T*_D_ (testing for the difference in cis-eQTL effect between CMC-brain and GTEx-blood) in each functional category (**Methods**). Error bars represent 95% confidence intervals around the estimates. The black dash line represents fold enrichment of l. Different colors in panels (A) and (B) correspond to 14 functional categories: TssA, active transcription start site; Prom, upstream/downstream TSS promoter; Tx, actively transcribed state; TxWk, weak transcription; TxEn, transcribed and regulatory Prom/Enh; EnhA, active enhancer; EnhW, weak enhancer; DNase, primary DNase; ZNF/Rpts, state associated with zinc finger protein genes; Het, constitutive heterochromatin; PromP, Poised promoter; PromBiv, bivalent regulatory states; ReprPC, repressed Polycomb states; and Quies, a quiescent state.

**Figure 3.**
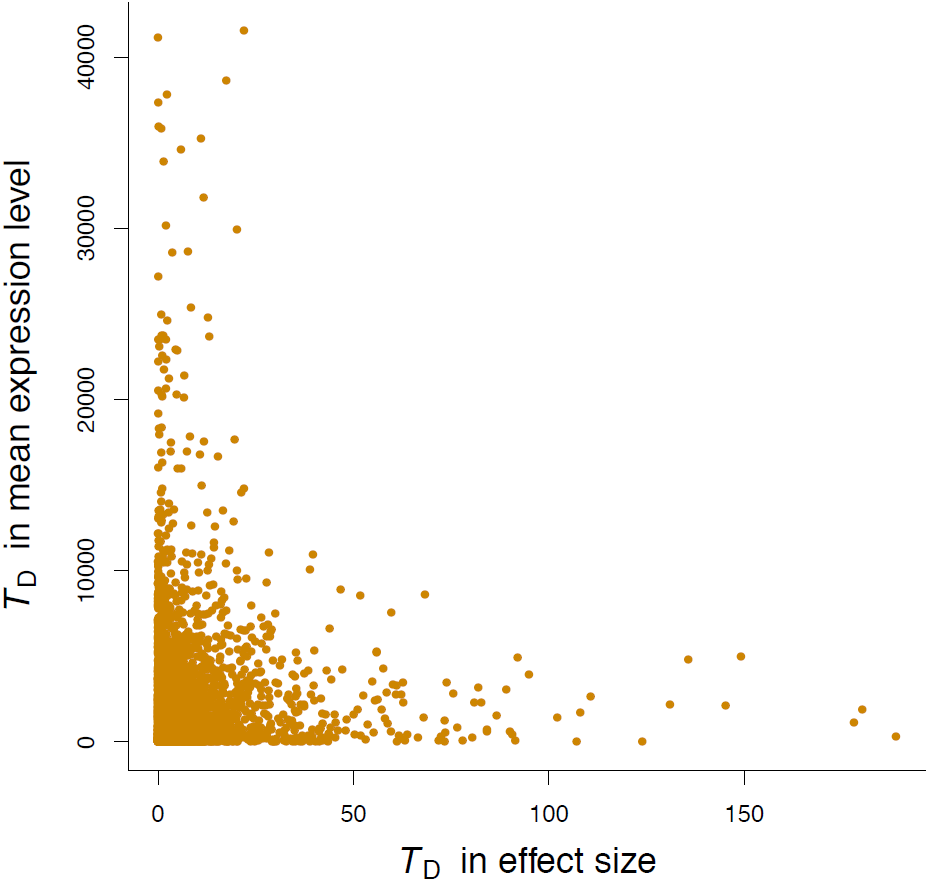
Relationship between the test-statistics for the difference (*T*_D_) in cis-eQTL effects and the *T*_D_ in mean expression level of the corresponding gene between GTEx-cerebellum and GTEx-blood for 3,569 genes. The 3,569 genes were ascertained with at least one cis-eQTL with *P*_eQTL_ < 5×10^−8^ in GTEx-muscle and expressed in GTEx-cerebellum and GTEx-blood (i.e. genes which have at least 10 samples with RPKM > 0.1 and raw read counts greater than 6). In this analysis, we used cis-eQTL effects in SD units and gene expression data in log(RPKM) units to avoid the confounding of the correlation by the mean-variance relationship in gene expression.

First, we selected the top associated cis-eQTLs at *P*_eQTL_ < 5×10^−8^ for 4,257 genes in GTEx-muscle and matched the selected genes with those in the other data sets (the number of matched genes ranged from 1,113 to 3,841) (**Supplementary Table 2**, i.e., up to 90%, with the lower numbers matched representing data sets with gene expression data for fewer genes). Note that all the matched genes were expressed in both tissues (i.e. genes which have at least 10 samples with reads per kilobase per million mapped reads (RPKM) > 0.1 and raw read counts greater than 6)^24^. Also note that our analysis below showed that there was no correlation between the test-statistics for tissue-specific gene expression and the test-statistics for tissue-specific SNP effects on gene expression, therefore selecting genes by cis-eQTL p-values would not bias mean gene expression in specific tissues. We used the Jackknife approach that removes one gene at a time to estimate the sampling variance of 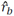 (**Methods**) assuming the estimated top cis-eQTL effects for different genes are independent. This assumption was approximately met given the small LD correlations among the 4,257 cis-eQTLs and the subtle difference between the mean Jackknife sampling variance and the observed sampling variance in simulation (**Supplementary Fig. 4**). The effect sizes of these cis-eQTLs were highly correlated between all the brain regions in GTEx after correcting for estimation errors, with a mean 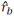 of 0.94 (SE = 0.004; **Fig. 1**). These estimates are higher than the Spearman correlations reported in a previous study^24^ because the Spearman correlation does not account for errors in the estimated SNP effects and therefore underestimate the correlation of true effects especially when the sample size is small. The two cerebellum measures (“brain cerebellar hemisphere” and “brain cerebellum”) appeared to be outliers. The correlation between “brain cerebellar hemisphere” and “brain cerebellum” was almost perfect (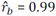 and SE = 0.002), but the correlations between these two regions and the other regions (mean 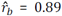 and SE = 0.006) were significantly smaller than the pairwise correlations between the other regions (mean 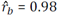 and SE = 0.003). We performed the same analysis in the Braineac data and observed similar results as above (**Supplementary Fig. 5**). The estimates of *r*_*b*_ between brain and blood in GTEx varied from 0.74 to 0.79 across different brain regions with a mean estimate of 0.77 (SE = 0.010). The estimates from ROSMAP were remarkably similar with those from CMC, providing an important replication of the result. The estimate of *r*_*b*_ between CMC (brain) and GTEx-blood was 0.74 (SE = 0.014), suggesting that the between-sample genetic heterogeneity is small, in line with the strong correlations between CMC and the brain regions of GTEx (mean 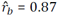 and SE = 0.010). The correlations related to Braineac were notably lower than those related to CMC (**Fig. 1**), which is likely due to the difference in transcriptomics technology between the two studies (microarray vs. RNA-Seq). The results were robust to scale transformation of the eQTL effects (**Supplementary Fig. 6**), the exclusion of cis-eQTLs in or near the promoter regions (**Supplementary Fig. 7**), the inclusion of secondary cis-eQTLs identified from a conditional analysis^25^ (**Supplementary Fig. 8**), or the adjustment of gene expression data for confounding (e.g. batch effects) predicted from the data (**Supplementary Fig. 9**). Second, we selected the top associated cis-eQTLs at *P*_eQTL_ < 5×10^−8^ from the CMC data, and found that the estimates of *r*_*b*_ among the brain regions and between brain and blood in GTEx remained largely unchanged (**Supplementary Fig. 10**), suggesting that our results are also robust to the ascertainment of the cis-eQTLs.

**Figure 4.**
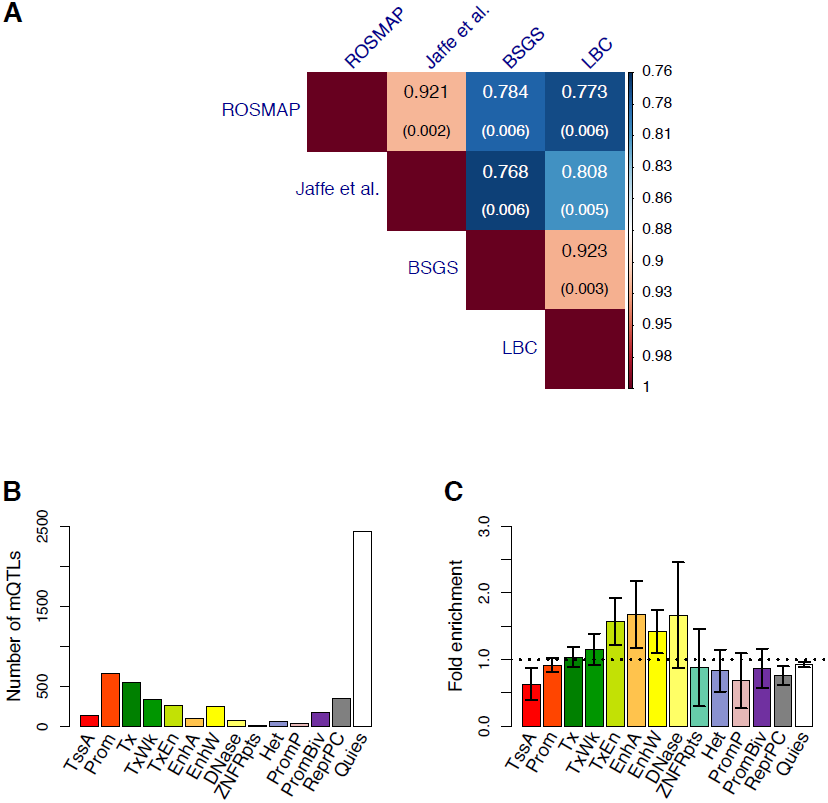
Similarity and difference in *cis*-mQTL effects between brain and blood. A) Estimated *r*_b_ for cis-mQTLs between brain and blood from 4 independent data sets. The cis-mQTLs (one for each DNAm probe) were selected at *P*_mQTL_ < 1×10^−10^ using data from the Hannon et al. study. Shown in each cell is the estimate of *r*_b_ with its standard error given in the parentheses (**Methods**). The distribution of cis-mQTLs across 14 functional categories derived from RMEC (**Methods**). Estimated enrichment of *T*_D_ (testing for the difference in cis-mQTL effect between Jaffe-brain and LBC-blood) in each functional category (**Methods**). Error bars represent 95% confidence intervals around the estimates. The black dash line represents the fold enrichment of l.

**Figure 5.**
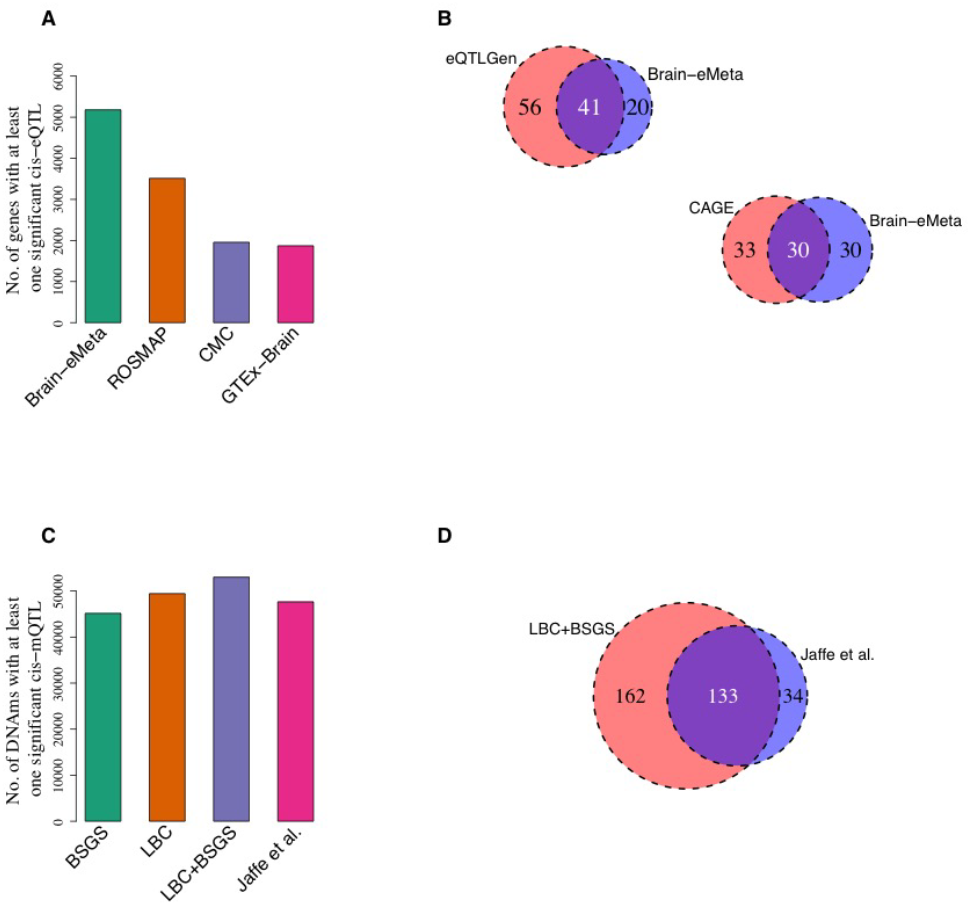
Identification of genes (DNAm sites) associated with 4 brain-related traits by an integrative analysis of GWAS data with eQTL (mQTL) data from brain and blood samples using the SMR method. The four brain-related traits are smoking, IQ, SCZ and EduYears. Panel A (C) shows the numbers of genes (DNAm sites) with a least one significant SNP at *P* < 5×10^−8^ in different data sets. Panel B (D) shows the numbers of genes (DNAm sites) associated with traits identified in different data sets. Sample sizes of the brain studies: GTEx-brain (*n* = ∼233), CMC (*n* = 467), ROSMAP (*n* = 494), Brain-eMeta (*n*_*eff*_ = ∼1,194) and Jaffe et al. (*n* = 526). Sample sizes of the blood studies: CAGE (*n* = 2,765), eQTLGen (*n* = 14,115), LBC+BSGS (*n* = l,980).

### cis-eQTLs with tissue-specific effects

The strong correlation of cis-eQTL effects between brain and blood (**Fig. 1**) does not preclude eQTLs with detectable difference in effect size between tissues. Of the 1,388 cis-eQTLs with *P*_eQTL_ < 5×10^−8^ in GTEx-muscle and available in CMC and GTEx-blood (**Supplementary Table 2**), 308 (22%) showed significant difference in effect size between CMC and GTEx-blood after Bonferroni correction for multiple testing (*P*_difference_ < 0.05/1,388) (**Methods**). It should be noted that the substantial proportion of eQTLs detected with significant between-tissue differences in effect sizes does not contradict the large estimate of *r*_*b*_ above (**Fig. 1**) because the power to detect a difference in effect size depends on sample size^13^ (**Supplementary Fig. 11**). Previous studies have indicated that functional variants (predicted by chromatin activity data) in enhancers are less likely to be shared across many tissues compared with those in promoters^26,27^, and that cell type-specific eQTLs are more dispersedly distributed around the transcription start site than eQTLs affected expression in multiple cell types^28,29^. These results seem to suggest that tissue-specific eQTLs are enriched in distal regulatory elements (i.e. enhancers) but the evidence are not direct. We computed the statistics to test for the between-tissue difference in effect size (denoted by *T*_D_) and tested the inflation (or deflation) of mean *T*_D_ of cis-eQTLs in the functional categories annotated by REMC (**Methods**). The result showed that although cis-eQTLs are enriched in genomic regions of active chromatin state (e.g. promoters and enhancers) and deflated in inactive regions, the mean *T*_D_ of cis-eQTLs between CMC and GTEx-blood was almost evenly distributed across all the functional categories with no evidence of inflation in the enhancer regions (**Fig. 2**). The result remained largely unchanged if we repeated the enrichment analysis based on *T*_D_ between GTEx-cerebellum and GTEx-blood (**Supplementary Fig. 12)**. There were some examples where the cis-eQTLs with tissue-specific effects in brain and blood were located in enhancers (**Supplementary Fig. 13**). These examples, however, were rare because only 14 of the 308 eQTLs with *P*_difference_ < 0.05/1,388 were located in enhancers and only 4 of the 14 enhancers appeared to be tissue specific. These results do not support the hypothesis that eQTLs with tissue-specific effects are more likely to be located in enhancers.

In addition, there are a large number of genes showing tissue-specific expression^11^. GWAS signals for a trait that are located in or near genes with tissue-specific expression are often seen as the evidence that the trait-associated genetic effects are enriched in particular tissues^30^. This implicitly assumes genetic variants with tissue-specific genetic effects on gene expression are co-located with genes with tissue-specific expression. We tested this hypothesis by examining the correlation between the test-statistic for the difference in cis-eQTL effect (in SD units) and the test-statistic for the difference in mean expression level of the corresponding gene (in log(RPKM) units) between GTEx-cerebellum and GTEx-blood for the 3,569 genes each with a cis-eQTL at *P*_eQTL_ < 5×10^−8^ in GTEx-muscle (**Supplementary Table 2**). It should be noted that the cis-eQTL effects were in SD units so that the correlation was not confounded by the mean-variance relationship in gene expression. We found that the correlation was marginal (*r* = 0.003) (**Fig. 3**), suggesting that genetic variants with tissue-specific genetic effects on gene expression (i.e. generating variation between people in a specific tissue) are not necessarily co-located with genes with tissue-specific gene expression (i.e. may be with genes that show similar levels of relative expression in other tissues). This is analogous to the observation that there is a large difference in the mean of height between men and women but the effect sizes of all autosomal SNPs on height in men are almost identical to those in women^31,32^. Therefore, the lack of enrichment of GWAS signals in or near genes over-or under-expressed in a tissue is not the evidence that the tissue is not relevant to the trait or disease. The lack of correlation between tissue-specific cis-QTL effect and tissue-specific expression level of the corresponding gene also means that the genes selected at *P*cis-eQTL < 5×10^−8^ in muscle for the *r*b analysis above were not necessarily enriched or depleted for tissue-specific expression. These results demonstrate the importance of generating tissue-specific eQTL data sets for integration with GWAS results and provide best bioinformatics functional annotation.

### Estimating the correlation of cis-mQTL effects between brain and blood

Having shown that cis-eQTL effects are highly correlated between brain and blood, we then turned to estimate the correlation of genetic effects on DNAm between the two tissues. To address this, we applied the *r*_*b*_ method developed above to mQTL data. We analyzed summary-level mQTL data from 5 studies based on the Illumina HumanMethylation450K array: fetal brain from Hannon et al. (*n* = 166)^33^, brain cortical region from ROSMAP (*n* = 468)^21^, frontal cortex region from Jaffe et al. (*n* = 526)^34^, and peripheral blood from McRae et al. (LBC: *n* = 1,366 and BSGS: *n* = 614)^35^ (**Supplementary Table 3**). All the mQTL effects are in SD units. We matched the SNPs in common across data sets, selected the top associated cis-mQTL at *P*_mQTL_ < 1×10^−10^ for 26,840 DNAm probes in the data from Hannon et al. (because only SNPs with *P*_mQTL_ < 1×10^−10^ are available in this data set) and matched the selected probes with those in the other data sets (the number of matched probes ranged from 4,892 to 6,561) (**Supplementary Table 4**). The correlation of cis-mQTL effects between two brain samples (ROSMAP and Jaffe et al.) was very strong (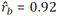 and SE = 0.002), similar to that between two blood samples (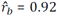 between BSGS and LBC with SE = 0.003) (**Fig. 4A**). It is of note that both estimates of *r*_*b*_ were smaller than unity, reflecting some degree of heterogeneity between studies. The mean brain-blood *r*_*b*_ estimate from two samples was 0.78 (SE = 0.006) (**Fig. 4A**), higher than that for cis-eQTLs (mean 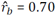 and SE = 0.015) shown above (**Fig. 1**). The result remained largely unchanged if the cis-mQTLs were selected at *P*_mQTL_ < 5×10^−8^ in the LBC data (**Supplementary Fig. 14**), again showing the robustness of our results to the choice of the reference tissue. In addition, of the 5,416 cis-mQTLs, 1,847 (34%) showed significantly different effects between brain (Jaffe et al.) and blood (LBC) after correcting for multiple testing (*P*_difference_ < 0.05/5,416). We then tested whether cis-mQTLs in any of the REMC functional categories tend to have higher *T*_D_ between brain and blood (see above). It seems that there were small but significant enrichments of *T*_D_ in enhancer regions (e.g. transcribed enhancer, active enhancer and weak enhancer) (**Fig. 4C**), and one of them survived multiple-testing correction (**Supplementary Table 5**).

### Meta-analysis of brain eQTL summary data from correlated samples

We know from the *r*_b_ analysis above that cis-eQTLs are almost perfectly correlated in different brain regions. We then sought to combine data from the brain regions to increase the power of detecting eQTLs for follow-up analysis (e.g. identification of putative functional genes for brain-related traits and diseases). However, if there is sample overlap between two tissues and the phenotypic correlation is nonzero, the estimation errors of the SNP effects from the two tissues will be correlated. We implemented in the SMR software tool a summary-data-based method, which only requires summary-level data in the cis-regions to account for sample overlaps, to <u>m</u>eta-analyze cis-<u>e</u>QTL data in <u>c</u>orrelated <u>s</u>amples (MeCS) (**Methods**). We showed by simulations that sample overlap could be estimated with high accuracy from the summary data of the null SNPs (e.g. *P*_eQTL_ > 0.01) in the cis-region using a simple correlation approach (**Supplementary Note**, **Supplementary Fig. 15B,** and **Supplementary Fig. 16A**), that the MeCS test-statistics were well calibrated under the null hypothesis (**Supplementary Fig. 15**), and that the MeCS estimates of meta-analysis effect sizes were well estimated under the alternative hypothesis (**Supplementary Fig. 16**). We compared MeCS to a univariate analysis of the mean expression phenotype across tissues, and found that the estimates of effect size and SE from the two approaches were highly consistent (**Supplementary Fig. 17**). It is of note that in comparison with the separate analysis in individual tissues, the gain of power for MeCS increased with the decrease of correlation in expression phenotype between tissues, more so for meta-analysis using individual-level data (**Supplementary Fig. 18**). The MeCS method has been implemented in the SMR software package (URLs). The method is general and can be applied to mQTL or even GWAS data.

We applied MeCS to data from 10 brain regions in GTEx (we referred to the meta-analyzed data as GTEx-brain hereafter). There were strong sample overlaps among the ten brain regions (mean overlap = 70.4%) and the mean correlation in expression level between pairwise brain regions across all the expressed genes was moderate (mean *r*_*p*_ = 0.33). The gain of power by the MeCS analysis was demonstrated by the observation that the mean *𝒳*^2^ statistic for cis-eQTLs (selected from GTEx-blood at *P*_eQTL_ < 5×10^−8^) in GTEx-brain was larger than that in any individual brain region (**Supplementary Fig. 19C**). The association test-statistic for a SNP written as *χ*^2^ = 1 + 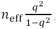, where *n* _*eff*_ is the effective sample size and *q*^2^ is the variance explained by a SNP^36^. We therefore can approximately estimate *n*_*eff*_ of GTEx-brain assuming constant mean *q*^2^ across brain regions (note that this assumption is justified by a mean *r*_*b*_ estimate of 0.94 between pairwise brain regions for cis-eQTL effects in SD units) (**Supplementary Note**). The estimate of *n*_*eff*_ of GTEx-brain was 233, approximately 2.6 times larger than the actual sample size of brain tissue in GTEx (mean *n* = ∼89 across 10 brain regions) (**Supplementary Fig. 19D**). To further increase the power of detecting brain eQTLs, we meta-analyzed GTEx-brain, CMC, and ROSMAP (referred to as Brain-eMeta hereafter). The gain of power is demonstrated by the increased number of genes with at least one cis-eQTL with *P*_eQTL_ < 5×10^−8^ in Brain-eMeta as compared with that in GTEx-brain, CMC, or ROSMAP (**Fig. 5A**).

### Identifying DNAm sites and genes associated with brain-related traits and diseases

With the Brain-eMeta eQTL data (*n*_*eff*_ = ∼1,194) obtained from the meta-analysis above, we applied the SMR approach^23,37^ to test for associations between gene expression levels with 4 brain related phenotypes, i.e. ever-smoked (smoking), fluid intelligence score (IQ), years of education (EduYears), and schizophrenia (SCZ). GWAS data were from published meta-analyses for EduYears and SCZ^38,39^, and from analyses of the full release of the UK Biobank data for smoking and IQ (**Methods and Supplementary Table 6**). LD data required for the HEIDI test^23^ were estimated from genotyped/imputed data of the Health and Retirement Study (HRS)^40^. LD *r*^2^ from HRS were strongly correlated with those from CMC (**Supplementary Fig. 20**), consistent with the observation from previous studies^25^. For power comparison, we included in the SMR analysis an additional set of blood eQTL data from a sample of 14,115 individuals from the eQTLGen Consortium. Only the genes with at least one cis-eQTL at *P*_eQTL_ < 5×10^−8^ (one of the basic assumptions of SMR) in both Brain-eMeta and eQTLGen were included. We further excluded genes in the major histocompatibility complex (MHC) region because of the complexity of this region, leaving 3,943 genes for analysis. We identified 61 genes associated with the traits using the brain eQTL data, 41 of which (67.2%) were in common with a larger set of genes (97) identified using the eQTLGen blood eQTL data (**Fig. 5B**). Despite the heterogeneity between the two eQTL data sets (Brain-eMeta was based on RNA-Seq and eQTLGen was based on microarray), the strong overlap between the two sets of results is consistent with the strong correlation of eQTL effects between brain and blood estimated above. For SCZ, 19 out of the 24 genes identified using brain eQTL data were replicated using blood eQTL data with an additional 27 genes identified only in the blood data because of its large sample size (**Supplementary Fig. 21**). We repeated the SMR analysis using blood eQTL data from the Consortium for the Architecture of Gene Expression (CAGE; *n* = 2,765)^9^ and observed similar result (**Fig. 5B**) although the power of CAGE was 1ower than that of eQTLGen (63 genes identified using CAGE versus 97 genes identified using eQTLGen).

We also performed the SMR analysis to detect associations between DNAm sites and the brain related phenotypes^17^ using brain mQTL data from Jaffe et al. (*n* = 526) and blood cis-mQTL data from a meta-analysis of LBC and BSGS (*n* = 1,980) (**Methods**). We only included in the analysis DNAm probes with at least one cis-mQTL with *P*_mQTL_ < 5×10^−8^ in both the brain and blood data sets. We identified 167 DNAm sites associated with the traits (*P*_SMR_ < 1.8×10^−6^) using the brain mQTL data, 133 of which (79.6%) were in common with the set of 295 DNAm sites identified using the blood mQTL data (**Fig. 5D** and **Supplementary Fig. 22**). The brain to blood “replication” rate slightly decreased when we rejected the associations with *P*_HEIDI_ < 0.05 (**Supplementary Fig. 23**), likely because of the HEIDI test being over-conservative especially as sample size increases^23^. These results further demonstrate the feasibility and gain of power of using the genetic effects on gene expression or DNAm estimated in blood to identify putative target genes and regulatory DNA elements for brain-related phenotypes.

## DISCUSSION

We introduced a summary-data-based method to estimate the correlation (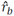) of genetic effects at the top associated cis-eQTLs/mQTLs between two tissues. Because the method accounts for estimation errors, 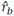 can be interpreted as an estimate of the correlation of true cis-eQTL effects between tissues, as demonstrated by simulations (**Supplementary Fig. 2**). We applied the method to summary-level eQTL data from GTEx and found that genetic effects on gene expression in the cis-regions were almost perfectly correlated between different brain regions (mean 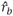 = 0.94 for cis-eQTLs), especially between the non-cerebellar regions (mean 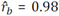 and SE = 0.003), in contrast to the modest phenotypic correlation in gene expression levels (mean *i*_*p*_ = 0.33). It is therefore sensible to run a meta-analysis of the cis-eQTL effects across brain regions to gain power of detecting eQTLs for the whole brain (**Supplementary Fig. 18**). This can be done even if the brain regions are from different samples. We developed the MeCS approach to meta-analyze cis-QTL data from independent or overlapping samples (only requires summary-level data of the SNPs in cis-regions to account for sample overlaps) and calibrated the method by simulations (**Supplementary Fig. 15 and Supplementary Fig. 16**). We applied MeCS to meta-analyze cis-eQTL summary data from the ten GTEx brain regions and demonstrated a ∼2.6 fold gain of power, on average, in comparison with any individual brain region. There is an existing method to conduct a joint analysis of summary statistics for multiple traits in overlapping samples (i.e. MTAG^41^). MTAG is a generalization of the inverse-variance-weighted meta-analysis. It relies on an estimate of sample overlap from bivariate LD score regression (LDSR)^42^ under a polygenic model. It is not applicable to our analysis which focused only on the SNPs in cis-regions.

We also found that the cis-eQTLs effects were highly correlated between brain and blood in GTEx (mean 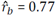 for cis-eQTLs), and the estimate only slightly decreased using data from different samples (mean 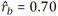). These estimates were significantly different from 1, suggesting there are real genetic differences between tissues. The genetic differences are partly due to cell-type specific genetic effects regardless whether cell composition have been included as covariates in the eQTL analysis or not. This is because adjusting for cell composition only removes the mean differences in gene expression level among cell types rather than cell-type specific genetic effects. On the other hand, however, the strong between-tissue correlation in cis-eQTL effects does not contradict the result that many genes showed differential expression across tissues because the difference in cis-eQTL effect is almost independent of the mean difference in gene expression level (**Fig. 3**). This is an important result and challenges a current dogma that focus interest on GWAS association results in genes that are differentially expressed in the tissue of most relevance to the disease. Our results reinforce the need to generate tissue-specific eQTL data sets to identify variants that generate variation between people in a specific tissue regardless of the relative expression level of the tissue.

Our results also provide some guidelines about the use of discovery-replication paradigm to compare eQTL effects between tissues (i.e. detecting eQTLs in one tissue at a stringent p-value threshold and replicating the effects in another tissue after correcting for multiple tests)^24,28^. Here, we often saw a low to moderate replication rate even if there is no genetic difference between the tissues. This is because the replication rate is a function of the sample size of the validation set (**Supplementary Fig. 11**) and the sample sizes of eQTL studies in non-blood tissues are often limited. If we apply the discovery-replication paradigm to the GTEx data, only ∼10.7% of eQTLs discovered in GTEx-muscle could be replicated in GTEx-hippocampus (although the estimates from the recent methods^43,44^ based on the discovery-replication paradigm were much higher) (**Supplementary Table 7**), which could potentially lead to a wrong conclusion that a large proportion of cis-eQTLs are tissue specific (note that the *r*_*b*_ estimate between the two tissues was 0.81). We therefore do not recommend the use of the discovery-replication paradigm to quantify the tissue-specific effects especially in small samples.

Data from genome annotation studies show that most enhancers are tissue specific^45^. In our study, we tested the difference in cis-eQTL effect between brain and blood, and did not observe an enrichment of the test-statistics (for tissue-specific cis-eQTL effects) in any of the functional annotation categories (**Fig. 2**). We performed a similar analysis for cis-mQTLs, and found a weak enrichment of the test-statistics (for tissue-specific cis-mQTL effects) in enhancer regions (**Fig. 4**). Because DNAm is an important epigenetic mechanism of regulating gene expression, we hypothesized that some of the tissue-specific cis-eQTL effects might be mediated through differentially methylated CpG sites.

We applied the SMR & HEIDI method to identify genes and DNAm sites that were associated with brain-related phenotypes through pleiotropy using summary data from GWAS and cis-eQTL/mQTL studies with large sample sizes (*n*_max_ = 453,693 for GWAS, *n*_max_ = 14,115 for eQTL and *n*_max_ = 1,980 for mQTL). We identified a number of genes and DNAm sites that showed pleiotropic associations with the phenotypes, consistent with a plausible model that the SNP effects on the phenotypes are caused by genetic regulation of the expression levels of the target genes and/or the methylation levels at the CpG sites. We repeated the analyses using eQTL and mQTL data from brain samples with much smaller sample sizes (*n*_max_ = 1,194 for eQTL and *n*_max_ = 526 for mQTL). Due to the lower power of the data sets, the number of genes or DNAm sites detected in the brain sample was much smaller than that using the blood sample (**Fig. 5**, **Supplementary Fig. 21, Supplementary Fig. 22, and Supplementary Fig. 23**), with at least 50% of genes (DNAm sites) in common between the two sets. These results provide strong justification of using blood samples to discover genes related to brain phenotypes and diseases. In practice, we recommend using a blood data set with large sample size for discovery, and an additional data set from brain for replication. This paradigm is certainly applicable to other tissues.

There are a few limitations in our study. First, our estimation of *r*_*b*_ are based on those genes which are expressed in both tissues. Genes that are only expressed in one tissue were not included in the estimation of *r*_b_. Therefore, the estimate of *r*_b_ needs to be interpreted with a restriction to genes expressed in both tissues. Although a quarter (4,257) of all genes were selected from GTEx-muscle, up to 90% of those selected genes were included in the *r*_b_ analysis. Second, we focused our analyses only on cis-eQTLs and cis-mQTLs because trans-eQTLs and trans-mQTLs data were not available in most data sets used in our study. Although most SNP-based heritability for gene expression levels are attributed to cis-eQTLs^9^, trans-eQTLs may also play an important role in regulating gene expression especially for tissue-specific effects^14^. The methods developed in this study can be applied to trans-eQTL/mQTL data with minimal modification. Because the variance explained by individual trans-eQTL/mQTL is small on average^9,35^, very large sample sizes (e.g. 10,000s) are required to detect trans-eQTLs to be useful for the SMR analysis^23^. Third, the *r*_*b*_ analysis was focused on the correlation at the top associated cis-eQTLs/mQTLs with relatively large effects (i.e. *P* < 5×10^−8^ in a reference tissue) because the SMR test only uses cis-eQTLs/mQTLs at *P* < 5×10^−8^. The estimate of *r*_*b*_ was slightly lower for cis-eQTLs/mQTLs selected at a less stringent threshold (**Supplementary Fig. 24**), consistent with the observation in simulation (**Supplementary Fig. 25**). However, this does not change our conclusion about the use of the top associated cis-eQTLs/mQTLs identified in a large blood sample to identify putative target genes for brain-related traits. Last but not least, the MeCS method requires the correlation of errors in the estimated SNP effects between two dependent samples (*θ*), which is estimated by a simple correlation approach at the null SNPs in the cis-region. This approach, however, is not applicable to ascertained eQTL or mQTL summary data by p-value. It will also be challenging to estimate *θ* if only a small number of cis-SNPs are available in the summary data. We therefore recommend eQTL and mQTL studies to make more cis-SNPs available without ascertainment (e.g. all the cis-SNPs in ±2Mb of the gene or DNAm). Despite these limitations, our findings shed light on the genetic architecture underlying the regulation of gene expression across tissues, and provide important guidance for studies in the future to identify functional genes for human complex traits.

## METHODS

### Summary data of cis-eQTL, cis-mQTL, and GWAS

All the analyses of eQTL/mQTL data were performed based on summary-level data from published studies. A summary description of all the data sets can be found in **Supplementary Table 1**, **Supplementary Table 3**, and **Supplementary Table 6**. All the samples were of European descent and the summary data available to us were derived from individual-level data that passed stringent quantify control (QC)^9,11,20,22,33-35,46^. The SNPs in all eQTL/mQTL data sets were from imputation of the genotyped data to the 1000 Genomes Project (1KGP) reference panels^47^, and only the common SNPs (MAF > 0.01) were included in analyses.

The eQTL summary-level data were from six published studies, i.e. the Genotype-Tissue Expression (GTEx)^11^ v6, the CommonMind Consortium (CMC)^20^, Religious Orders Study and Memory and Aging Project (ROSMAP)^21^, the Brain eQTL Almanac project (Braineac)^22^, the Architecture of Gene Expression (CAGE)^9^ and eQTLGen. In GTEx, ROSMAP, and CMC, gene expression levels were measured by RNA-Seq. Genes in GTEx and ROSMAP were annotated by GENCODE^48^ v19 and v14 respectively, and genes in CMC were annotated by Ensembl. We accessed the GTEx eQTL summary statistics of ∼9.3 million SNPs for ∼32,000 genes in 44 tissues (including 10 brain regions) through GTEx portal (**URLs**). The sample sizes of different tissues in GTEx ranged from 70 to 361 with an average of 160. We accessed the CMC summary data from Synapse (accession: syn2759792). The CMC eQTL summary statistics (ascertained at FDR<0.2 in the public domain) of ∼1.1 million SNPs for 14,366 genes were derived from individual-level data in dorsolateral prefrontal cortex of 467 subjects, 209 of which were schizophrenia patients. We accessed the ROSMAP eQTL summary statistics of ∼6.4 million SNPs for 12,979 genes, which were derived from individual-level data in dorsolateral prefrontal cortex of 494 subjects. We accessed the Braineac eQTL summary statistics of ∼6.2 million SNPs for 25,490 genes, which were derived from data in 10 brain regions of 134 subjects free of neurodegenerative disorders^22^. The gene expression levels in Braineac were measured by Affymetrix Human Exon 1.0 ST Arrays. For blood eQTL data, we used eQTL summary data from CAGE^9^ (38,624 gene expression probes and ∼8 million SNPs on 2,765 subjects) and eQTLGen (44,556 gene expression probes and ∼10 million SNPs on 14,115 subjects). Gene expression levels in CAGE and eQTLGen were measured by Illumina gene expression arrays. We mapped the probes to genes based on the annotations provided by Illumina. The eQTL summary data available in GTEx, CAGE, and eQTLGen were from previous analyses of standardized gene expression levels with mean 0 and variance 1 whereas expression levels in the other data sets (i.e. CMC, ROSMAP, and Braineac) were not standardized, resulting in differences in the units of eQTL effects among data sets. To harmonize the units across data sets, we re-scaled the effect size and standard error (SE) of each eQTL in the CMC, ROSMAP, and Braineac based on the z-statistic, allele frequency and sample size using the method described in Zhu et al.^23^ so that the eQTL effects in all data sets can be interpreted in standard deviation (SD) units.

mQTL summary statistics were from 5 data sets: brain cortical region from ROSMAP study (*n*_ind_ = 468, *n*_probe_ = 420,103, *n*_snp_ = 5 million)^21^; fetal brain from Hannon et al. (*n*_ind_ = 166, *n*_probe_ = 26,840, *n*_snp_ = 0.3 million)^33^; frontal cortex region from Jaffe et al. (*n*_ind_ = 526, *n*_probe_ = 138,917, *n*_snp_ = l.5 million)^34^; and peripheral blood from McRae et al.^35^ (Lothian Birth Cohorts^49^ (LBC): *n*_ind_ = 1,366 and Brisbane Systems Genetics Study^50^ (BSGS): *n*_ind_ = 614). DNAm levels in all these five studies were based on the Illumina HumanMethylation450K array. We performed a meta-analysis of LBC and BSGS, resulting in 397,621 DNAm probes and ∼7.7 million SNPs. The DNAm levels of all the five studies were not standardized. We computed the effect size and SE of each mQTL from their z-statistics using the method described in Zhu et al.^23^.

We included in the analysis 4 brain-related complex traits, i.e. ever-smoked (smoking), fluid intelligence score (IQ), years of education (EduYears), and schizophrenia (SCZ). GWAS summary statistics for EduYears (*n* = 293,723) and SCZ (36,989 cases and 113,075 controls) were from the latest meta-analyses^38,39^, and summary data for smoking (*n* = 453,693) and IQ (*n* = 146,819) were from GWAS analyses of the latest release of the UK Biobank (UKB) data^51^. Quality control and imputation of the UKB data have been detailed elsewhere^51^. We used 456,426 individuals of European descent and 7,288,503 common SNPs (MAF > 0.01) imputed from the Haplotype Reference Consortium (HRC)^52^ reference panel in the analysis. IQ was measured by 13 fluid intelligence questions and detailed description of the measurement can be found in http://biobank.ctsu.ox.ac.uk/. We adjusted IQ (*n* = 146,819) by age and sex, and standardized the adjusted phenotype by rank-based inverse-normal transformation. The GWAS analyses were performed in BOLT-LMM^53^ using all 7.3 million SNPs with a subset of 0.7 million SNPs in common with HapMap3^54^ used to control for population structure and polygenic effects. We used self-reported “ever smoked” as a dichotomous phenotype for smoking (208,988 cases and 244,705 controls). We analyzed the data in BOLT-LMM based a linear model with age and sex fitted as covariates, and transformed the effect size of each SNP on the observed 0-1 scale to odds ratio (OR) using LMOR (http://cnsgenomics.com/shiny/LMOR/).

### Correlation of cis-eQTL effects between tissues

Let 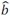 be the estimated effect size of the top associated cis-eQTL for a gene. We can model 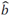 as

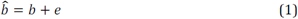

where *b* is true effect size and *e* is the estimation error. We assume that *b* and *e* are random variables when interrogated across genes, i.e. *b*∼*N (0, var (b))* and *e*∼*N (0, var (e))*. The covariance of the estimated cis-eQTL effects between tissues *i* and *j* across a number of genes can be partitioned into the covariance of the true cis-eQTL effects and the covariance of estimation errors due to sample overlap, i.e.

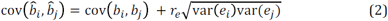

where *var (e*_*i*_) and *var (e*_*j*_) are the variance of the estimation error in tissues *i* and *j* respectively, and *r*_*e*_ is the correlation of estimation errors across genes between two tissues, i.e. *r*_*e*_ *=* cor*(e*_*i*_*, e*_*j*_). We know from Bulik-Sullivan et al.^42^ and Zhu et al.^55^ that *r*_*e*_ *≈ r*_*p*_*ρ*, where 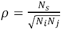 measures the sample overlap with *N*_*i*_ and *N*_*j*_ being the sample sizes in tissues *i* and *j* respectively, *N*_*s*_ the number of overlapping individuals, and *r*_*p*_ is the correlation of gene expression levels between two tissues in the overlapping sample. If *i = j*, then *r*_*e*_ *=* 1 and 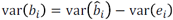, where *var (b*_*i*_) is the variation of true cis-eQTL effects across genes. We therefore can estimate the correlation of true cis-eQTL effect sizes across genes as

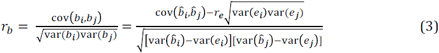

where 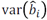, 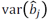 and 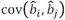 can be observed from the eQTL summary data, and *var (e)* is the variation of the estimation errors in estimated cis-eQTL effects across genes. The reported SE of the estimated eQTL effect is an estimate of the standard deviation of the estimation error. We therefore can estimate *var (e)* by the mean SE squared across genes. We know from equation (2) that if *b*_*i*_ *= b*_*j*_ *= 0*, 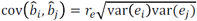 Hence,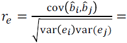 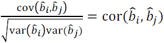 for null SNPs. In practice, we estimated *r* for each “null” SNP (*P*_eQTL_ > 0.01) in the cis-region by a simple correlation approach and took the average across SNPs.

The sampling variance of 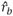 is computed via Jackknife approach leaving one gene out at a time.

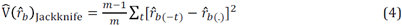

where 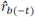 is the estimate with the *t*-th gene left out and 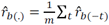 The method is derived based on eQTL data but can be applied to data from genetic studies of different types of molecular phenotypes (e.g. DNAm and histone modification).

### Enrichment of cis-eQTLs with tissue-specific effects in functional annotations

We used chromatin state data from 23 blood samples (T-cell, B-cell and Hematopoietic stem cells) and 10 brain samples generated by the NIH Roadmap Epigenomics Mapping Consortium (REMC)^19^. There were 25 chromatin states predicted by ChromHMM^56^ based on the imputed data of 12 histone-modification marks^19^. We classified the 25 chromatin states into 14 main functional categories by combining functionally relevant annotations. We tested the difference in eQTL effect for a gene between two tissues (*i* and *j*) using the method below. Let

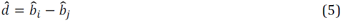

The sampling variance of 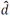 can be written as

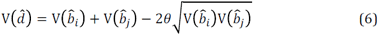

where 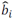 and 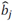 are the estimated effect sizes of the top associated cis-eQTL for a gene in two tissues, 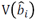 and 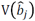 are the sampling variance for 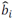 and 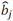, respectively, and *θ* is sampling correlation between 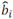 and 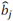 for the gene. 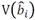 and 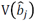 can be estimated by the squared SE for 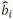 and 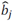, and *θ* can be estimated from all the “null” SNPs (e.g. *P*_eQTL_ > 0.01) in the cis-region for each gene using the simple correlation approach described above. The significance of 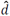 can therefore be assessed by a Wald test, i.e., 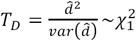.

To test the enrichment of *T*_D_ statistics in functional annotations, we allocated the cis-eQTLs to the 14 functional categories described above by physical position, and calculate the mean *T*_D_ of each category. We assessed the enrichment using the inflation factor 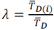, where 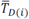 is the mean *T*_D_ of the cis-eQTLs in a category *i*, and 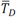 is the mean *T*_D_ of all the cis-eQTLs. We then used the Jackknife approach (leaving one gene out at one time) described above to compute the variability of *λ.* Note that although we described the enrichment test method above based on cis-eQTLs, the method can be applied to data from genetic studies of different types of molecular phenotypes (e.g. DNAm and histone modification).

## Meta-analysis of cis-eQTL data from correlated samples

We know from equation (1) that the estimated effect of a cis-eQTL for a gene can be partitioned into two components, i.e. the true effect size (*b*) and the estimation error (e). For multiple tissues, the joint distribution of the estimates can be written as

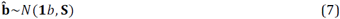

where 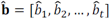, **S** is the sampling (co)variance matrix with 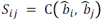, which can be estimated by *θS*_*i*_*S*_*j*_ when *i* ≠ *j*, where *θ* is sampling correlation between 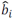 and 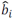 for the gene, and *S*_*i*_ and *S*_*j*_ are the SEs of 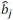 and 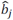 respectively. If *i* = *j*, then *θ = 1* and 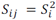 In practice, we can use the simple correlation approach described above to estimate *θ* from all the “null” SNPs (e.g. *P*_eQTL_ > 0.01) in the cis-region for each gene. Similar to the summary data based meta-analysis methods that account for correlated estimation errors^57,58^, we can estimate combined effect as

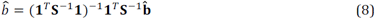

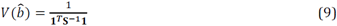

The significance of 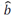 can be assessed by a Wald test, i.e. 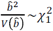.

## SUPPLEMENTAL INFORMATION

Supplemental data include Supplementary Note, 25 supplemental figures and 7 supplemental tables.

## WEB RESOURCES

MeCS, http://cnsgenomics.com/software/smr/#MeCS

SMR, http://cnsgenomics.com/software/smr

GTEx Portal, http://www.gtexportal.org/

CMC, https://www.synapse.org/CMC www.synapse.org/CMC Braineac, http://www.braineac.org/

Brain-eMeta eQTL summary data will be available at the SMR website when the manuscript is formally accepted (http://cnsgenomics.com/software/smr/#Download).

## ACKNOWLEDGEMENTS

This research was supported by the Australian Research Council (DP160101343, DP160101056, DP160103860 and DP160102400), the Australian National Health and Medical Research Council (1113400, 1107258, 1083656, 1078037 and 1078901), the US National Institutes of Health (GM099568, GM075091 and AG042568), and the Sylvia & Charles Viertel Charitable Foundation. This study makes use of data from dbGaP (accessions: phs000428.v1.p1 and phs000424.v6.p1), UK Biobank Resource (application number: 12514), UK10K project and <u>CommonMind Consortium</u>. A full list of acknowledgements to these data sets can be found in **Supplementary Note**. The members of the eQTLGen Consortium are (in alphabetical order): Mawussé Agbessi, Habibul Ahsan, Isabel Alves, Anand Andiappan, Philip Awadalla, Alexis Battle, Frank Beutner, Marc Jan Bonder, Dorret Boomsma, Mark Christiansen, Annique Claringbould, Patrick Deelen, Tõnu Esko, Marie-Julie Favé, Lude Franke, Timothy Frayling, Sina Gharib, Gregory Gibson, Gibran Hemani, Rick Jansen, Mika Kähönen, Anette Kalnapenkis, Silva Kasela, Johannes Kettunen, Yungil Kim, Holger Kirsten, Peter Kovacs, Knut Krohn, Jaanika Kronberg-Guzman, Viktorija Kukushkina, Zoltan Kutalik, Bernett Lee, Terho Lehtimäki, Markus Loeffler, Urko M. Marigorta, Andres Metspalu, Lili Milani, Martina Müller-Nurasyid, Matthias Nauck, Michel Nivard, Brenda Penninx, Markus Perola, Natalia Pervjakova, Brandon Pierce, Joseph Powell, Holger Prokisch, Bruce Psaty, Olli Raitakari, Susan Ring, Samuli Ripatti, Olaf Rotzschke, Sina Ruëger, Ashis Saha, Markus Scholz, Katharina Schramm, Ilkka Seppälä, Michael Stumvoll, Patrick Sullivan, Alexander Teumer, Joachim Thiery, Lin Tong, Anke Tönjes, Jenny van Dongen, Joyce van Meurs, Joost Verlouw, Peter Visscher, Uwe Völker, Urmo Võsa, Hanieh Yaghootkar, Jian Yang, Biao Zeng, Futao Zhang.

